# Defining *Target Population of Environments* to Enviromics Studies Using R-based GIS Tools

**DOI:** 10.1101/2024.09.25.614957

**Authors:** Demila D. M. Cruz, Alexandre B. Heinemann, Gustavo E. Marcatti, Rafael T. Resende

## Abstract

We propose an R-based function that facilitates the definition of TPE (Target Population of Environments) as GIS polygons for enviromics studies in plant breeding. By adjusting parameters such as pixel size, buffers, and concavity, this function enhances envirotypic-based G×E interaction analysis and provides a flexible tool to optimize environmental and spatial assessments.

## 1. Introduction

The Target Population of Environments (TPE) serves as the foundational framework for identifying the locations where plant varieties or cultivars are grown, guiding the selection of genotypes with enhanced adaptation and performance (Cooper et al., 2023; Crespo-Herrera et al., 2021). In the context of Enviromics (Resende et al., 2021; Resende et al., 2024a), the TPE accommodates high-resolution environmental data on a Geographic Information Systems (GIS) scale with detailed phenotypic data. This integration enables a more precise characterization of environmental conditions, facilitating the model prediction of genotype performance in different scenarios and the adjustment of selection to the specific conditions of each location within the TPE (Piepho, 2022). Thus, the TPE evolves, incorporating big data, making breeding programs more efficient by directing cultivars to specific environments and maximizing genetic gains in the TPE (Costa-Neto and Fritsche-Neto, 2021; Xu et al., 2022).

Clipping the TPE in a GIS context involves challenges that require the integration of detailed spatial and environmental data (Resende et al., 2024a). It is essential to consider geographic and climatic variability, as well as cover distinct soil classes and agricultural practices within each target region; hence, there is a need to use GIS tools to map and analyze these aspects. In simplified terms, TPE refers to an area—whether continuous or discontinuous—where a genotype or group of genotypes exhibit their performances. Therefore, TPE must ensure that selected environments are representative, facilitating the development of new varieties by integrating geospatial data with information on genotype × environment (G×E) interactions (Cooper et al., 2023). Moreover, the spatial and temporal dynamics of environmental conditions, such as climate change and the expansion of cultivation areas, must be considered. To capture all these conditions, concepts such as the surrounding polygon from Multi-Environmental Trials (MET) can be employed, with either a Concave or Convex Hull (Gombin et al., 2020) providing precise delineation of the TPE.

The creation of a computational tool based on R and GIS is a pathway to assist plant breeders in defining the TPE with precision. By integrating phenotypic data from MET or commercial fields (on-farm trials) with geospatial information, this tool maps and visualizes the spatial distribution of cultivation environments in detail. This facilitates the identification of environmental patterns and variability within the target area, ensuring that the TPE represents the actual cultivation conditions (Resende et al., 2024b). The use of GIS allows these data to be cross-referenced with climatic, edaphic, and management information (Pebesma and Bivand, 2023), providing an integrated view of the factors influencing genotype performance. This work introduces the R function ’**TPEmap**’, which adds buffers and adjusts concavity parameters to clip a precise TPE, applicable to any crop species. Here, we illustrate a practical application of this approach using a common bean (*Phaseolus vulgaris* L.) dataset from the Embrapa Arroz & Feijão Breeding Program, as detailed in Heinemann et al. (2022).

## 2. Methods

The concepts of TPE, Mega-environments, and Breeding Zones are complementary, though often confused. Here we will describe a function to define geographic TPE polygons, called ’**TPEmap**’, which will delineate the entire area where experimental or on-farm trial data points serve as reference. The *premise* is: if some crop is being breeding tested or growing at a location, it is part of the target-environment, and the surrounding areas, which are also of interest, are included to capture environmental variability. Mega-environments complement this by grouping broad regions with similar agro-climatic conditions, helping to select genotypes adapted to these larger regions (Gauch & Zobel, 1997). Breeding Zones can further refine TPEs, identifying specific sub-regions where G×E interaction is minimized, and fine-tuning the selection process for local conditions (Resende et al., 2024). While TPEs broadly define target environments (Crespo-Herrera et al., 2021), Mega-environments and Breeding Zones optimize selection across different scales, enhancing precision and efficiency (Cooper et al., 2023).

### 2.1 Geospatial Data Processing and Analysis

The geospatial data processing began with loading the geographic coordinates of the MET and on-farm trial points, using the ’**sf**’ library in R for manipulating shapefiles (Pebesma and Bivand, 2023). The coordinates were loaded and transformed into the coordinate reference system (CRS) WGS 84 (EPSG:4326), ensuring spatial data compatibility. The ’generate_coordinates’ function was created to test different scenarios, allowing the simulation of any number of points within a given area. Here, we used Brazil as a reference, but any area across the globe can be set.

After processing and preparing the geospatial data, the next step involved applying buffers to the geographic coordinates (see the Trial Buffers in Figure 1), representing phenotypic data collection points, such as breeding experiments or on-farm data. Using the ’**st_buffer**’ function from the ’**sf**’ package, buffers were applied at tuning scales of kilometers around each point, creating polygons that were then merged using ’**st_union**’ to form a single aggregated area of influence.

**Figure 1.**
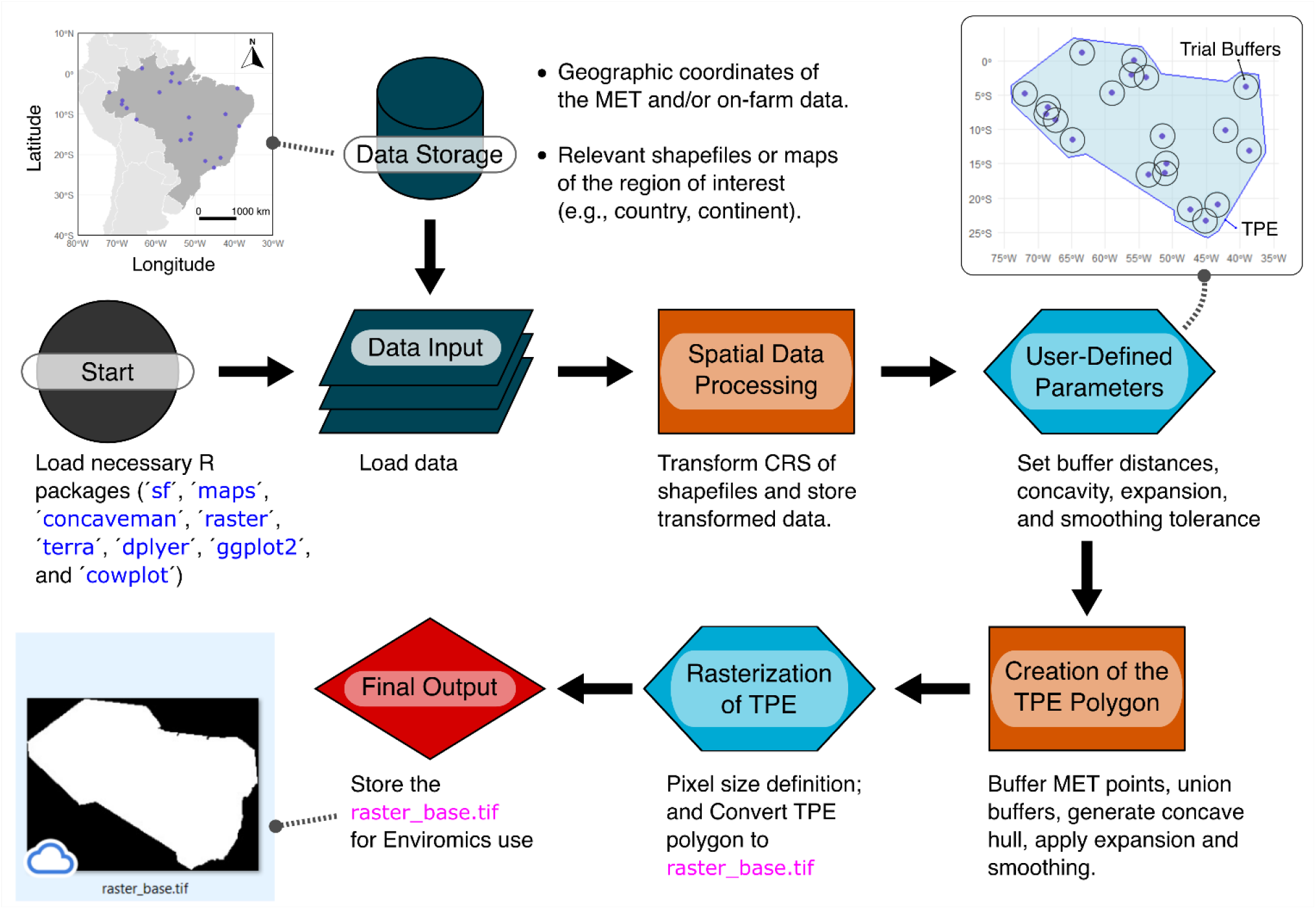
GIS Flowchart of the steps for constructing a TPE (Target Population of Environments) using the GIS-based ’TPEmap’ function (Geographic Information Systems). The process includes the loading and transformation of geospatial data, the definition of user-defined parameters (buffers, concavity, smoothing), the creation of the TPE polygon, and the final rasterization. The result is the raster_base.tif file, used for Enviromics analyses, integrating data from MET (Multi-Environmental Trials) or on-farm trials with environmental information.

### 2.2 Building the TPE Polygon Using a Concavity Algorithm

The construction of the polygon that defines the TPE was carried out using the ’**concaveman**’ algorithm in R (Gombin et al., 2020), which enables the creation of a concave polygon adaptable to the geospatial points of interest. After generating the buffers around the trial points, these polygons were merged and converted into a set of points to form the basis of the concave polygon. The ’**TPEmap**’ function allows the user to adjust parameters such as buffer distance, concavity, and length threshold, customizing the polygon according to the specific needs of the study. The ’**generate_coordinates**’ and ’**TPEmap**’ functions, with instructions for different applications, are available on GitHub at: https://github.com/enviromics/TPE-mapping.

Thru the TPE polygon generation, an additional buffer can be applied, allowing the user to expand the final area to include all desired influence zones. The whole TPE polygon can be smoothed to remove irregularities and small protrusions, with the user having the flexibility to adjust the smoothing tolerance through the ’**st_simplify**’ function. The breeder may also choose to define an external TPE without including trials containing phenotypic data, aiming for the prospecting of predictive results; however, this approach cannot be validated using appropriate cross-validation schemes (Resende et al., 2024b).

The final polygon is then converted into a raster, with pixel size defined according to the G×E study’s specifications. The resulting file, raster_base.tif, is prepared for use in Enviromics analyses, integrating environmental data with the geographical location of the trials, ensuring that the TPE represents the environmental conditions of the regions of interest. Refer to Resende et al., (2024b) for some potential Enviromics results that can be achieved using the base raster. It is important to note that increasing pixel resolution (i.e., reducing its granularity), while desirable, will result in a quadratic increase in the total number of pixels within the TPE, leading to the need for greater physical computational memory.

## 3. The ’TPEmap’ Function Arguments

The ’**TPEmap**’ function was developed to assist in the creation of a TPE for plant breeding, based on geospatial data from MET or on-farm trials. The arguments for this function include essential adjustments (Figure 1, ‘User-Defined Parameters’), such as buffers, polygon concavity, and pixel size for rasterization, which will be explained in detail below:

- ’**coordinates**’: A data frame with X (longitude) and Y (latitude) columns, representing the geographic coordinates of MET or on-farm trial points.
- ’**point_buffer**’: A numeric value that defines the buffer distance to be applied around each point (in kilometers). This argument allows the user to adjust the size of the area of influence around the points.
- ’**concavity**’: A numeric value that defines the degree of polygon concavity. Lower values create a more detailed polygon, while higher values (up to infinity) resulting in a Convex Hull.
- ’**length_threshold**’: A numeric value that defines the edge length threshold in the concavity algorithm. Edge segments with lengths below this value are not considered for additional detail. This argument allows the user to control the level of detail in the polygon.
- ’**expansion_buffer**’: A numeric value that defines the additional buffer distance to be applied after the concave polygon is generated, expanding the final TPE (in kilometers).
- ’**simplify_tolerance**’: A numeric value that defines the simplification tolerance of the final polygon. This option allows the user to smooth the polygon, removing excessive detail and small irregularities.
- ’**pixel_size**’: A numeric value that defines the pixel size for rasterizing the TPE. This argument is used in the conversion of the final polygon to a raster.

## 4. Features of the ’TPEmap’ Function

Adapting the ’**generate_coordinates**’ function, and to provide a more realistic example of TPE creation, we used simulated geographic coordinates for 100 geographic points (e.g., experiments or plantings), matched for suitability with soybean cultivation as suggested by Silva et al. (2021). In these simulations, protected/natural reserve areas were excluded. Figure 2 illustrates different possible configurations using ’**TPEmap**’, allowing the user to define buffer sizes, adjust the concavity of the generated polygon (Concave Hull), as well as set the length threshold (’**length_threshold**’) and simplification tolerance (’**simplify**’).

**Figure 2.**
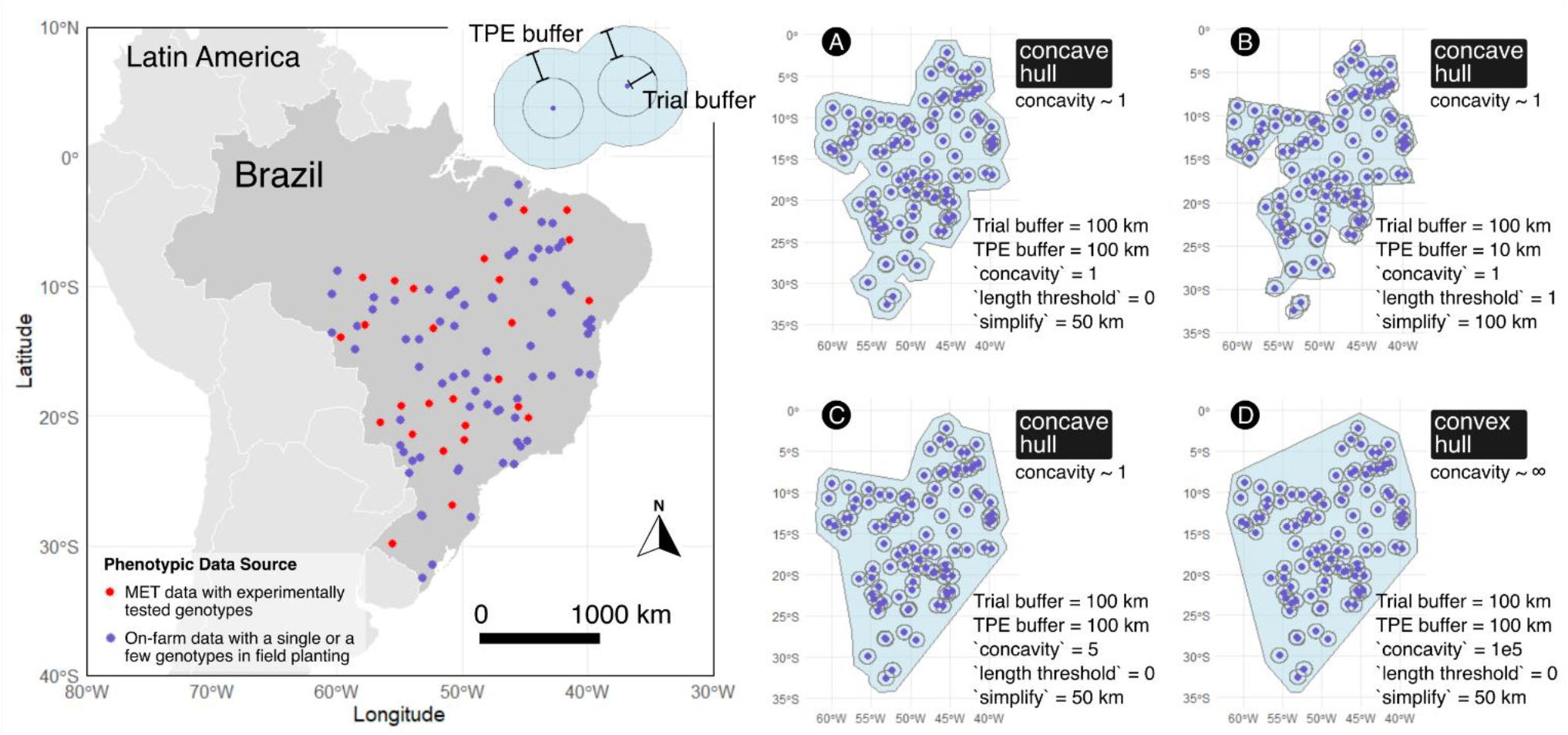
Definition of the TPE (Target Population of Environments) polygon using different concavity and buffer parameters. (**A**) TPE polygon generated with a concavity of ∼1, a trial buffer of 100 km, a TPE buffer of 100 km, and a simplification of 50 km. This area includes 100 geographic points, of which 26 are breeding experiments (the red ones). (**B**) TPE polygon with a concavity of ∼1, a trial buffer of 100 km, a TPE buffer of 10 km, and a simplification of 100 km. (**C**) TPE polygon with a concavity of ∼1, a trial buffer of 100 km, a TPE buffer of 100 km, and a concavity adjusted to 5. (**D**) TPE polygon generated as a Convex Hull (concavity ∼∞), with a trial buffer of 100 km and a TPE buffer of 100 km. The different concavity and buffer configurations directly influence the shape and fit of the polygon, reflecting the areas of interest for Enviromics analysis.

Concavity and length threshold parameters are fundamental tunes in TPE clipping, allowing the user to generate polygons that range from highly detailed shapes (Figure 2A and 2C) to simpler and more convex shapes (Figure 2D). The function also enables the visualization of different adjustments, such as varying the concavity from 1 to 5, and even using a Convex Hull (infinite concavity, we adopted 1e5 in this context), highlighting the different ways to encapsulate trial points.

## 5. Application to Common Bean MET Data set

In the process of defining the TPE for common bean (*Phaseolus vulgaris* L.), we used MET coordinates data from Embrapa Arroz & Feijão, covering three different crop season types: Rainfed (Figure 3-A), Dryland (Figure 3-B), and Winter (Figure 3-C) (Heinemann et al., 2022). Among these, there are 423 trials conducted at 71 unique geographic locations between 2011 and 2023, across up to three different crop seasons, depending on the regionalization of production areas (Figure 3-D), specifically: 26 were conducted in the Rainfed season, 12 in rotations of Dryland and Rainfed, 9 in Rainfed and Winter, 9 exclusively in Winter, 7 exclusively in Dryland, 5 in Dryland and Winter, and 3 locations covered all three seasons. For reference, 87 commercial cultivars are present in these tests, with 52 of them being “Carioca” type and 35 “Preto” type.

**Figure 3.**
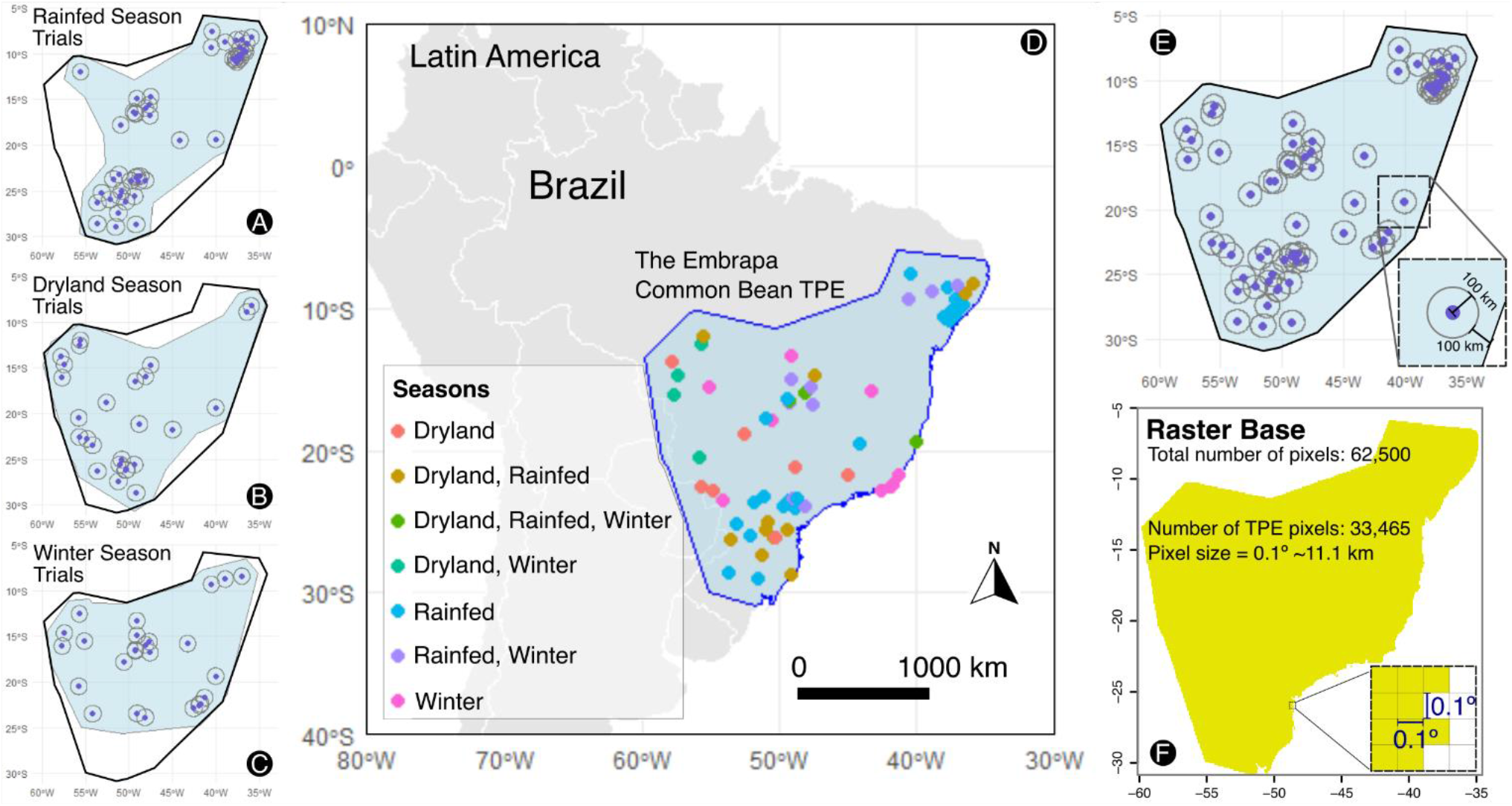
Definition of the TPE for common bean based on trial types (seasons). (**A**) TPE polygon for rainfed season trials after applying a trial buffer. (**B**) TPE polygon for dryland season trials. (**C**) TPE polygon for winter season trials. (**D**) Final TPE polygon combining all common bean season trials across different seasons, marked by various colors representing dryland, rainfed, and winter seasons. (**E**) Illustration of the combined trial buffers (incorporating the three types of seasons) and the TPE, showing the buffer distances applied around the trial points. (**F**) Rasterized TPE base with a pixel size of 0.1° (∼11.1 km), displaying the total number of pixels and the number of pixels within the TPE.

Figure 3-E illustrates how the surrounding polygons (Concave Hull) were generated using a concavity of 2 and a ’**length_threshold**’ of 10, with the inclusion of trial buffers (’**point_buffer**’) of 100 km around each point. After applying the buffers, the TPE polygon was expanded with an additional 100 km buffer, totaling 200 km, and smoothed with a simplification tolerance of 50 km. Identifying and mapping representative cultivation areas, considering the varied environmental conditions faced by common bean in Brazil, is critical, especially given the country’s bioclimatic diversity, as highlighted in the study by Elias et al. (2021).

Considering the different responses of common bean to climatic variations in each crop season is key to adjusting the TPE to the actual cultivation conditions in Brazil (Heinemann et al., 2022). The results show a wide dispersion of experimental points across the Brazilian territory, covering various climatic zones and soil types. The application of the ’**TPEmap**’ function allows the construction of a TPE polygon that integrates data from the three seasons, providing a representation of common bean cultivation areas. The overlay of different trials, as shown in Figure 3D, shows that considering multiple seasons is an operational alternative for defining the TPE, ensuring broad environmental variability. However, a TPE can be broad, season-specific, or based on other breeding program stratification strategies.

Considering modern envirotyping techniques, as highlighted by Xu et al. (2020) and Resende et al. (2024a), the rasterized base with refined pixels can then be used to acquire the environmental covariates needed for Enviromics studies and G×E interactions at different spatial scales (Resende et al., 2021). Envirotyping data can be obtained from platforms such as WorldClim, Planet, NASAPower, ERA5, SRTM, MODIS, and SoilGrids (see more details in Resende et al., 2024a), or through the elegant ’**EnvRtype**’ package, which facilitates access to NASAPOWER data (Costa-Neto et al., 2021). It is essential to balance the resolution of each platform, as the ’**raster_base**’ may have a high level of refinement, while data sources may offer coarser pixels. When converting the TPE to a raster base (Figure 3F), a pixel size of 0.1° (∼11.1 km) was used, resulting in 33,465 pixels within the TPE. On the other hand, refining to 0.01° increases the number of pixels in the TPE to 3,347,285, which will pose computational challenges for downloading envirotyping data.

The results from Resende et al. (2024a) highlight the importance of environmental variables such as soil, radiation, and temperature in predicting genotypic performance. The choice of pixel resolution in defining the TPE influences the quality of the representation, with smaller pixels offering greater detail but requiring a balance between precision and computational efficiency, as emphasized by Piepho (2022). This work underscores the precise definition of TPEs using GIS tools, enabling breeders to consider geographic variation and optimize G×E interaction analysis in envirotyping studies, enhancing the adaptation and performance of varieties across different environments.

